# Protein phosphatase 1 regulates huntingtin exon 1 aggregation and toxicity

**DOI:** 10.1101/135954

**Authors:** Joana Branco-Santos, Federico Herrera, Gonçalo M. Poças, Yolanda Pires-Afonso, Flaviano Giorgini, Pedro M. Domingos, Tiago F. Outeiro

## Abstract

Huntington’s disease (HD) is neurodegenerative disorder caused by a polyglutamine expansion in the N-terminal region of the huntingtin protein (N17). Here, we analysed the relative contribution of each phosphorylatable residue in the N17 region (T3, S13 and S16) towards huntingtin exon 1 (HTTex1) oligomerization, aggregation and toxicity in human cells and *Drosophila* neurons. We used bimolecular fluorescence complementation (BiFC) to show that expression of single phosphomimic mutations completely abolished HTTex1 aggregation in human cells. In *Drosophila*, Mimicking phosphorylation at T3 decreased HTTex1 aggregation both in larvae and adult flies. Interestingly, pharmacological or genetic inhibition of protein phosphatase 1 (PP1) prevented HTTex1 aggregation in both human cells and *Drosophila* while increasing neurotoxicity in flies. Our findings suggest that PP1 modulates HTTex1 aggregation by regulating phosphorylation on T3. In summary, our study suggests that modulation of HTTex1 single phosphorylation events by PP1 could constitute an efficient and direct molecular target for therapeutic interventions in HD.

## Introduction

Huntington’s disease (HD) is characterized by the loss of medium spiny neurons in the striatum. The main histopathological hallmark of HD is the misfolding and subsequent intracellular aggregation of a mutant form of huntingtin (HTT) [1]. HTT is a very large protein (∼350 kDa), but expression of exon 1 is sufficient to produce HD-like features in various cellular and animal models [2–4]. HTT exon 1 (HTTex1) contains a polyglutamine (polyQ) tract that, in normal conditions, is constituted by 6 to 35 glutamine residues. An expansion of the polyQ tract beyond 35 glutamines induces the misfolding and aggregation of mutant HTT and causes HD [5,6]. Mutant HTT with longer polyQ expansions is more prone to aggregate, and leads to earlier onset of the disease [2,7,8].

The polyQ tract is preceded by an N-terminal sequence of 17 amino acids (N17 domain) that is highly conserved, suggesting that this domain plays an important role in the function of HTT [9–15]. The N17 domain plays a key role in the aggregation pathway of HTT, where the protein associates first into alpha-helical oligomers and acts as a seed to concentrate and facilitate the formation of larger aggregates and fibrils [16–20]. Deletion or posttranslational modifications such as phosphorylation, ubiquitination and SUMOylation in the N17 produce striking effects in the stability and aggregation of HTT, as well as in cell viability [21–34]. The N17 domain has 3 phosphorylatable amino acid residues – threonine at position 3 (T3), and serine residues at positions 13 and 16 (S13 and S16). Constitutive phosphorylation of T3 enhances mutant HTT aggregation, but its role in HTT toxicity remains unclear [29]. Previous studies have focused on double S13/S16 phosphorylation [30,31,35–38], despite the fact that double S13/S16 phosphorylation is less frequent than single T3 or S13 phosphorylation [39,40], and that overexpression of particular kinases is required in order to achieve double S13/S16 phosphorylation [30].

Single phosphorylation and phosphomimetic modifications in S13 or S16 modulates the formation of HTT fibrils and reduces the oligomerization rate in cell-free systems, just as the double S13/S16 phosphorylation does, both *in vitro* and *in vivo* [26,28,30,31]. Casein Kinase 2 (CK2) inhibitors reduce S13/S16 phosphorylation and enhance toxicity [37], while GM1 ganglioside induces S13/S16 phosphorylation and restores motor and molecular deficits in HD mice [36]. IKK, a kinase involved in inflammatory responses, also regulates T3 and S13/S16 phosphorylation and modulates HTT aggregation [30,37,39,41]. While several protein phosphatases control HTT dephosphorylation beyond exon 1 and modulate its toxicity [42–46], it is not known which protein phosphatases, if any, regulate phosphorylation of the N17 domain.

Here, we elucidate the contribution of single N17 phosphorylation events towards HTTex1 oligomerization, aggregation and toxicity. We screened a collection of protein phosphatase chemical inhibitors to identify modulators of HTTex1 oligomerization and aggregation in human cells. Inhibition of PP1 prevents HTTex1 aggregation but not oligomerization in human cells. In addition, downregulation of PP1 in *Drosophila* neurons reduces HTTex1 aggregation and increases its toxicity. In total, our findings point to a critical role of T3 phosphorylation in HTTex1 aggregation and support the targeting of PP1 for therapeutic interventions in HD.

## Results

### Single N17 phosphomutants modulate HTTex1 aggregation in human cells

In order to investigate the contribution of each phosphorylatable residue within the N17 region (T3, S13 and S16) towards HTTex1 aggregation, we used the BiFC system for the visualization of both oligomeric species and inclusion bodies of HTTex1 in living cells, which we have previously described [47–49]. In this system, wild type (19Q) or disease-causing (97Q) HTTex1 are fused to non-fluorescent halves of the Venus fluorescent protein (Fig 1A). In this system, upon dimerization of HTTex1 fragments, the Venus halves are brought together and reconstitute the functional fluorophore. Therefore, fluorescence is proportional to the extent of HTTex1 dimerization/oligomerization. We introduced point mutations in each of the phosphorylatable residues within the N17, changing these amino acids to either alanine (A mutants, which cannot be phosphorylated - phosphoresistant) or to aspartic acid (D mutants, which mimic the phosphorylated state - phosphomimic). Importantly, these phosphomutants behave like phosphorylated peptides in terms of their aggregation in cell-free systems and primary neuronal cultures [26,28].

**Figure 1.**
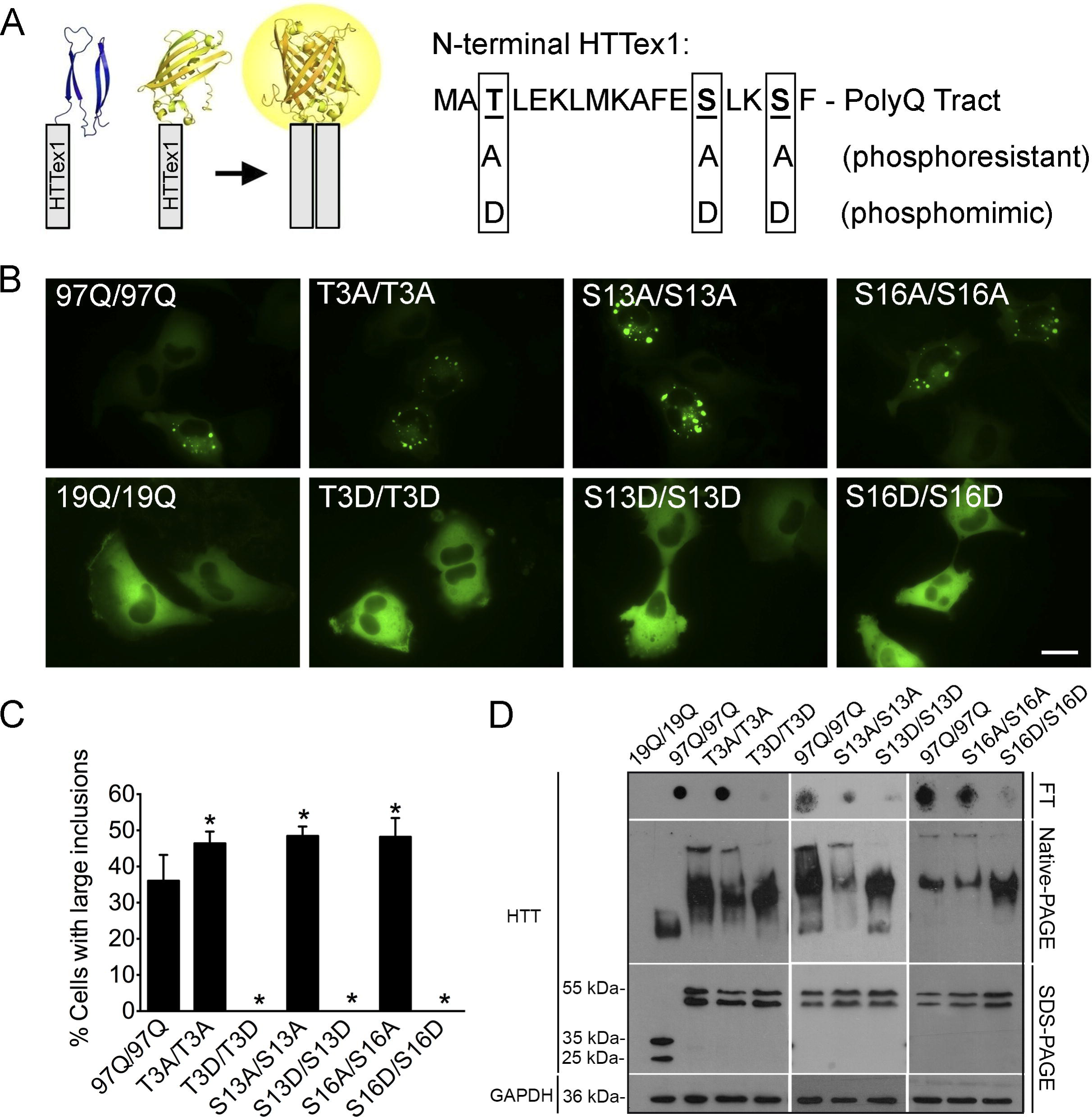
Single N17 phosphomimic mutations modulate 97QHTTex1 aggregation but not oligomerization in human H4 glioma cells. (A) HTTex1 (gray bars) was fused to two non-fluorescent halves of the Venus protein. When HTTex1 dimerizes, Venus recovers fluorescence. The three N17 phosphorylatable residues were mutated to mimic phosphorylation or dephosphorylation. (B) Cells transfected with phosphoresistant BiFC pairs resemble the non-mutated 97QHTTex1 phenotype in terms of aggregation, while phosphomimic pairs showed a total absence of aggregates. Scale bar, 20 μm. (C) Quantitative analyses of microscopy pictures. Data are represented as average ± SD of at least 3 independent experiments. *, significant versus 97Q/97Q, p<0.05 (one-way ANOVA). (D) Filter trap assays (FT) were consistent with microscopy results. Phosphomimic pairs did not produce insoluble aggregates (FT), but showed similar levels of oligomeric species (Native-PAGE) comparing to non-mutated 97QHTTex1. Expression levels of each pair were evaluated by SDS-PAGE.

We had previously shown that 97QHTTex1-Venus BiFC pairs oligomerize and aggregate more readily than wild-type 19QHTTex1-Venus BiFC pairs [49]. Strikingly, all single phosphomimic mutations of 97QHTTex1-Venus BiFC constructs (T3D, S13D, S16D) completely abolished the formation of inclusion bodies in human cells (Fig 1B and C)while the phosphoresistant mutants (T3A, S13A, S16A) behaved similarly to non-mutated 97QHTTex1-Venus BiFC pairs (Fig 1B and C and S1 Fig). These phenotypes were further confirmed by filter trap assays, where non-mutated 97QHTTex1 and phosphoresistant pairs appear as large SDS-insoluble aggregates, as opposed to wild-type 19QHTTex1 and phosphomimic pairs. (Fig 1D, filter trap (FT)). No differences in oligomerization/fluorescence levels were observed between the phosphomutants, and non-mutated 97QHTTex1, as determined by flow cytometry (S2 Fig). Native-PAGE analyses confirmed that all phosphomutants formed oligomeric species (Fig 1D). These results are consistent with a recent report where expanded HTTex1 was shown to exist as tetramers but not monomers [50].

Due to its dynamic nature, phosphorylation usually affects only a fraction of the total pool of any given protein. Thus, we hypothesized that subpopulations or pools of 97QHTTex1 with different extents of phosphorylation may co-exist in cells. In order to determine whether 97QHTTex1 fragments with different phosphorylation status interacted in living cells, we took advantage of the unique features of our BiFC system to screen all possible pairwise combinations of phosphomutants and non-mutated 97QHTTex1 control (Table 1). Combinations of phosphomimic with non-mutated 97QHTTex1 oligomerized to the same extent as the non-mutated 97QHTTex1 pair, as determined by flow cytometry (S3 Fig). However, phosphomimic pairs did not form inclusions, independently of the mutated residue (Table 1). Combinations of phosphomimics with non-mutated 97QHTTex1 (Table 1 and Fig 2A and B) or phosphoresistant mutants (Table 1) resulted in intermediate aggregation phenotypes, depending on which residue was mutated. Combinations including T3D, S13D or S16D resulted in 0 %, 14.1-22.0 % or 19.8-29.1 % of cells with inclusions, respectively (Table 1), which is significantly less than the percentage of cells with inclusions observed with the non-mutated 97QHTTex1 pair (36.1%). In contrast, phosphoresistant pairs generally resulted in an increased percentage of cells displaying inclusions (43.5-48.5% cells with inclusions), with the exception of the S13A/S16A combination (35.3%) (Table 1).

**Table 1.** Percentage of cells containing HTT inclusions (average ± SEM) when co-transfected with different combinations of HTTex1-, Syn- or Tau-Venus BiFC plasmids.

|  |  | Venus 2 |  |  |  |  |  |  |  |  |
| --- | --- | --- | --- | --- | --- | --- | --- | --- | --- | --- |
|  |  | T3D | S13D | S16D | T3A | S13A | S16A | 97Q | Syn | Tau |
| Venus 1 | T3D | 0 | 0 | 0 | 0 | 0 | 0 | 0 | 0 | 15.5±2.2 |
| | S13D | | 0 | 0 | 22.0±3.4 | 17.1±2.3 | 14.5±1.2 | 14.1±1.3 | 5.6±1.7 | 15.7±2.2 |
| | S16D | | | 0 | 27.5±1.5 | 29.1±3.6 | 19.8±3.0 | 22.9±1.0 | 10.0±4.8 | 16.5±1.7 |
| | T3A | | | | 46.4±1.3 | 43.5±3.6 | 45.9±2.5 | 34.5±2.2 | N/D | N/D |
| | S13A | | | | | 48.5±1.3 | 35.3±2.4 | 37.8±1.9 | N/D | N/D |
| | S16A | | | | | | 48.2±2.6 | 36.6±6.1 | N/D | N/D |
| | 97Q | | | | | | | 36.1±2.2 | 39.3±3.9 | 16.2±1.3 |

N/D, not determined.

**Figure 2.**
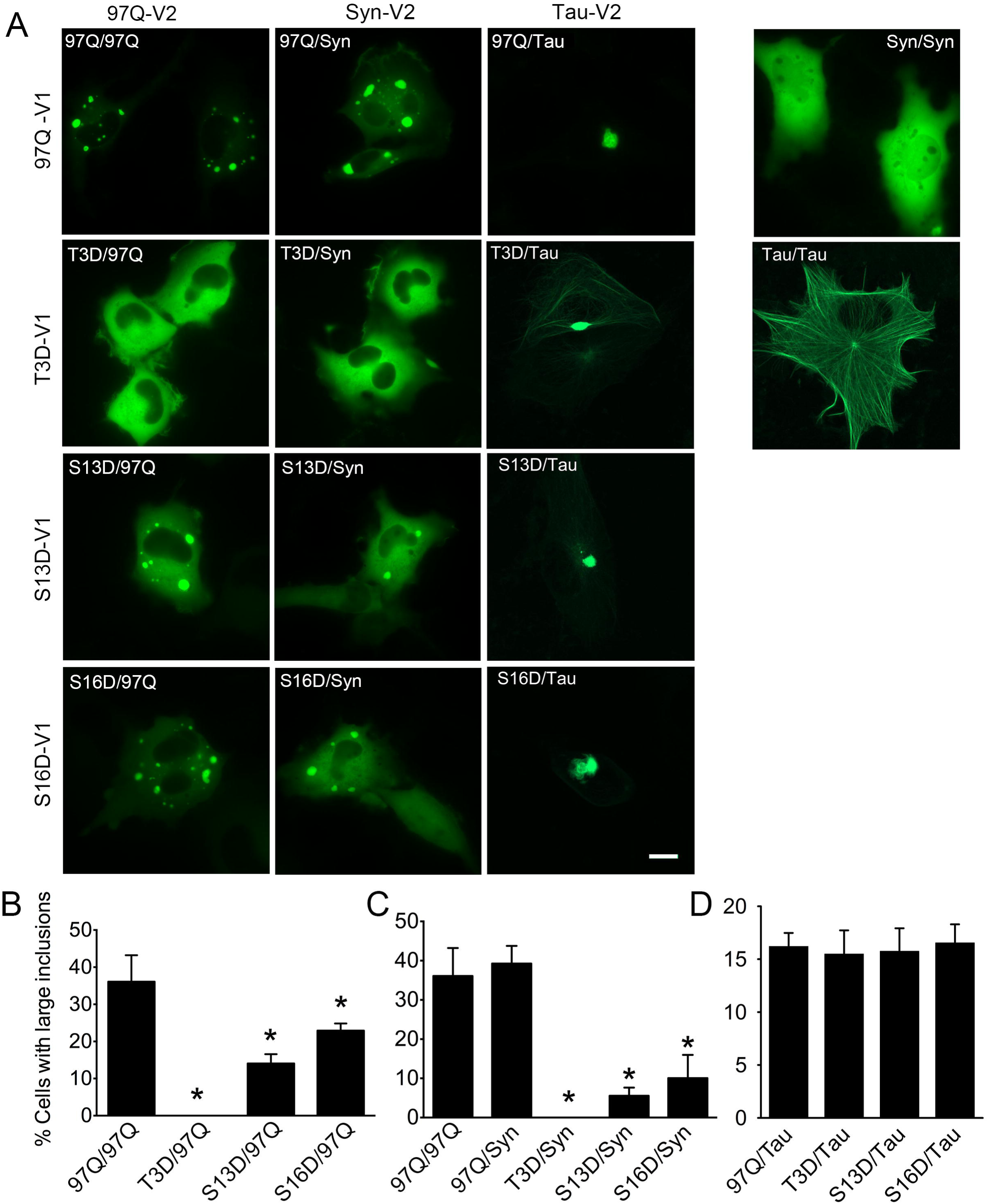
Effect of phosphomimic mutants on 97QHTTex1 aggregation is protein-specific. (A-D) Combinations of 97QHTTex1-Venus 1 BiFC constructs with synuclein- or tau-Venus 2 BiFC constructs revealed different co-aggregation responses to phosphomimic constructs. 97QHTTex1 (B) and synuclein (C) showed similar patterns of aggregation in the presence of single phosphomimic mutations, with T3D being the most restrictive for decreasing aggregation. (D) Tau co-aggregated with 97QHTTex1 regardless of the N17 mutation. Data information: Data are shown as average ± SD of at least 3 independent experiments. *, significant versus 97Q/97Q (B) or 97Q/Syn (C), p<0.05 (one-way ANOVA). Scale bar, 20 μm.

Importantly, the effect of phosphomimic mutants on 97QHTTex1 aggregation appears to be protein-specific (Fig 2). We and others have recently shown that HTTex1 co-aggregates with alpha-synuclein (aSyn) and Tau, and that these interactions change the aggregation profile of HTTex1 [47,48,51,52]. Combinations of phosphomimic 97QHTTex1 with aSyn BiFC constructs resulted in a residue-dependent reduction in the percentage of cells with inclusions, with the T3D/Syn showing no inclusions (Fig 2A, C). On the other hand, combinations of Tau-Venus plasmids showed the same aggregation pattern independently of the mutated residue (Fig 2A, D). These observations further support that single N17 phosphomutants modulate HTTex1 aggregation, and that the T3D is the most restrictive modification in preventing HTTex1 aggregation.

### Increased dynamics of 97QHTTex1 inclusions containing S13 or S16 phosphomimic mutants

We next analysed the dynamics of 97QHTTex1 inclusions using fluorescence recovery after photobleaching (FRAP). We observed a faster fluorescence recovery in inclusions formed by pairs containing phosphomimic mutants (S13D or S16D) and non-mutated 97QHTTex1 in comparison to inclusions formed exclusively by non-mutated 97QHTTex1 (Fig 3A and 3B, S1-S3 Videos). The phosphoresistant (T3A or S13A)/97QHTTex1 pairs, but not the S16A/97QHTTex1, presented slower fluorescence recovery in inclusions (Fig 3C). Phosphomimic-containing inclusions often had less defined boundaries, appearing rather continuous with the cytosolic fluorescence, consistent with areas of more dynamic exchange of HTT protein. Inclusions formed by the S13D/97QHTTex1 combination (Fig 3D and S4 Video) showed a fluid-like behaviour as opposed to the more rigid behaviour of the inclusions formed by combinations containing phosphoresistant mutants (S5 Video) or non-mutated 97QHTTex1 BiFC pairs (S6 Video).

**Figure 3.**
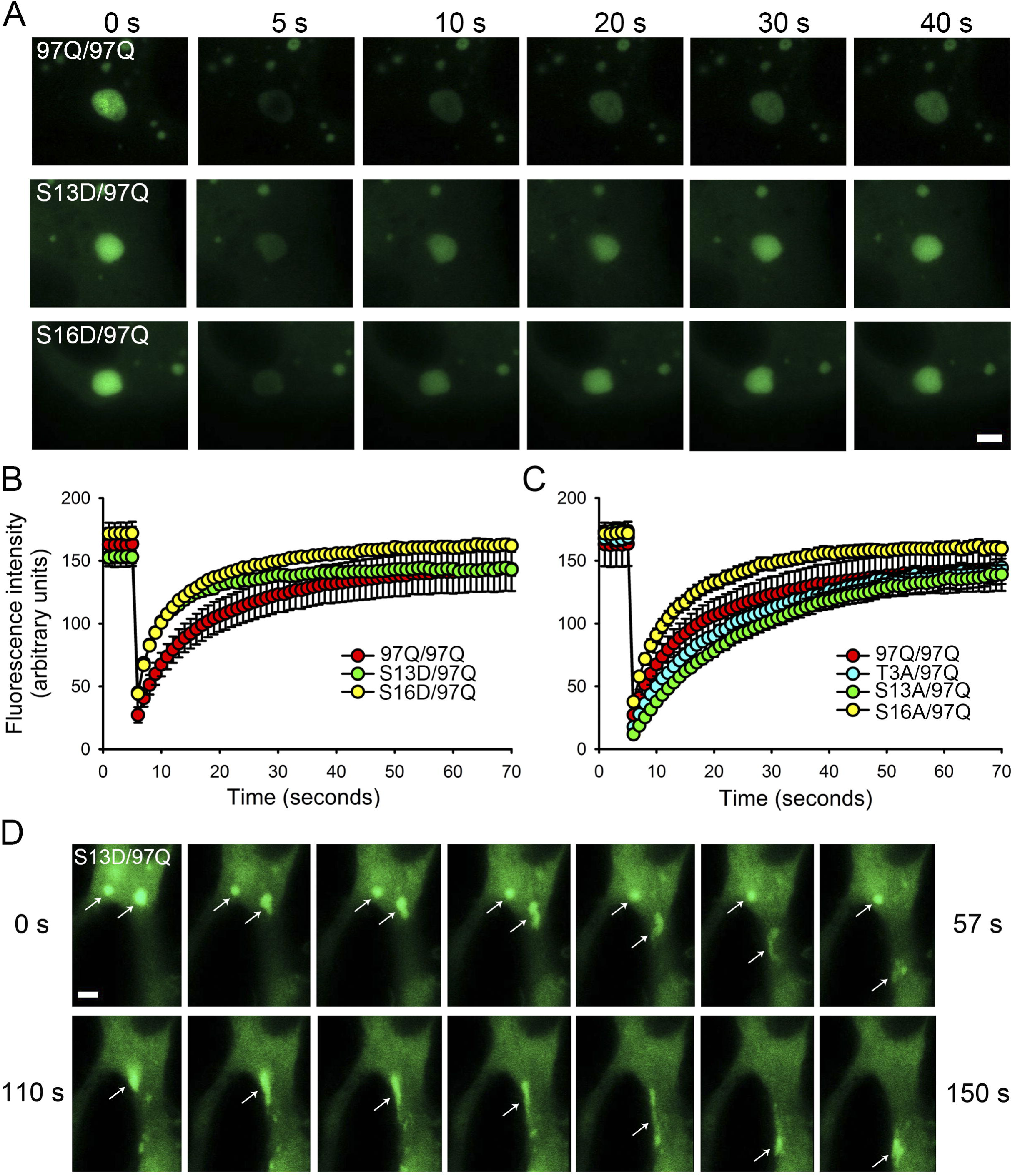
Phosphomimic-containing inclusions of 97QHTTex1 are more dynamic. (A) Time lapse of FRAP experiments on aggregates containing non-mutated 97QHTTex1 alone or in combination with phosphomimic mutant in H4 glioma cells. T3D mutants produced no aggregates under any circumstance. (B) Aggregates containing S13D or S16D mutants recovered significantly faster than the non-mutated 97QHTTex1. (C) Aggregates combining non-mutated 97QHTTex1 with T3A or S13A recovered more slowly than aggregates made only of non-mutated 97QHTTex1, with the exception of the S16A/97Q combination. (D) Time lapse of aggregates formed by non-mutated 97QHTTex1 and S13D (S4 Video). Aggregates containing phosphomimic versions are highly dynamic and flexible, and move (arrow indicates moving aggregate in each frame) through the cell with more freedom than non-mutated 97QHTTex1 aggregates (S1 Video). Data information: Scale bars, 20 μm.

Overall, the results suggest that single phosphorylation events within N17 domain prevent HTTex1 aggregation but not its oligomerization, and that phosphorylation at the T3 residue might play a critical role in modulating HTTex1 aggregation. In addition, our observations support the idea that inclusions might be composed of non-phosphorylated HTT or mixtures of non-phosphorylated and phosphorylated pools of molecules, consistent with data indicating that disease-causing HTT is hypophosphorylated in the N17 region [29,37].

### Protein phosphatases regulate HTTex1 aggregation in human cells

The results above indicate that single N17 phosphorylation can modulate HTTex1 aggregation, which is consistent with the idea that N17 phosphorylation might be enhanced by the activation of kinases [30,37,39] or, alternatively, by the inhibition of phosphatases. In order to identify protein phosphatases that might mediate HTTex1 dephosphorylation, we screened a library of 33 phosphatase chemical inhibitors for their effect on HTTex1 aggregation and oligomerization, using our 97QHTTex1-Venus BiFC system (Fig 4, and S1 Table). We found that inhibitors of PP1/PP2A, CD45, and Cdc25 prevented 97QHTTex1 aggregation as determined by filter trap assay (Fig 4A). These results were confirmed by fluorescence microscopy, where we observed that inhibitors for those phosphatases significantly decreased the percentage of cells with inclusions (Fig 4B). Interestingly, the decrease in oligomerization upon treatment with these phosphatase inhibitors was much less dramatic than the reduction observed in aggregation levels (B01, B02, B07, B08 and C04). Although a slight reduction in fluorescence/oligomerization was observed for 28 out of the 33 inhibitors assayed, the few exceptions where a striking decrease was observed (C06, D01 and D03) were due to cytotoxicity (S1 Table). Importantly, cells treated with CD45 inhibitors showed reduced levels of HTTex1 expression and increased toxicity (S1 Table), and were therefore excluded from the study.

**Figure 4.**
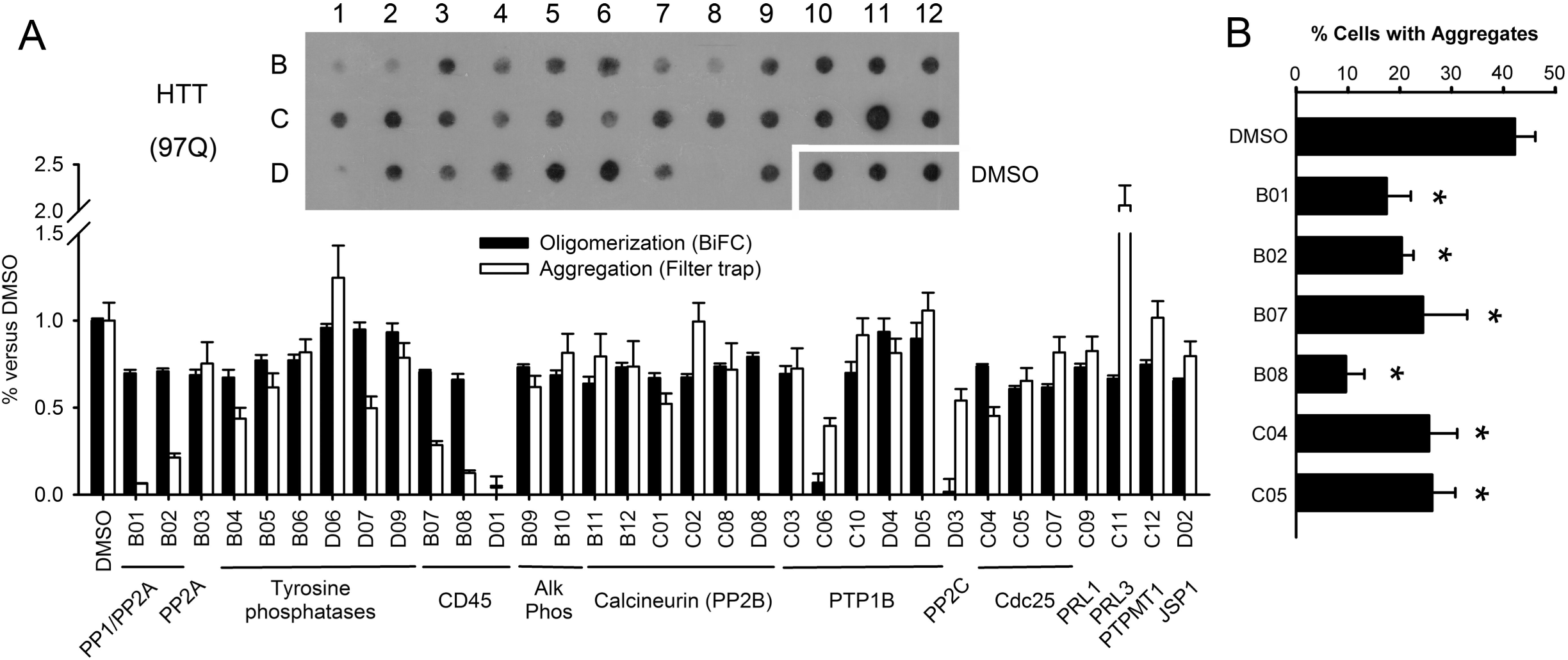
Protein phosphatases regulate HTTex1 aggregation in mammalian cells. (A) Representative filter trap of protein extracts from cells transfected with non-mutated 97QHTTex1 BiFC constructs and treated with chemical inhibitors of protein phosphatases. The graph shows oligomerization levels as determined by flow cytometry (black bars) and aggregation levels as determined by optic density of filter trap dots (white bars). The names of phosphatase inhibitors (Enzo Life Sciences) and the concentrations used are described in S1 Table. (B) Selected inhibitors PP1/PP2A (B01 and B02) and Cdc25 (C04 and C05) also reduced the percentage of H4 cells with large HTTex1 inclusions as determined by quantitative analyses of microscopy pictures. Data information: Data are all average ± SD of at least 3 independent experiments. *, significant versus control treated with DMSO, p<0.05 (one-way ANOVA).

### Single N17 phosphomutants modulate HTTex1 aggregation in *Drosophila*

To further investigate the role of N17 phosphorylation on HTTex1 aggregation, we expressed single phosphoresistant or phosphomimic versions of 97QHTTex1 in *Drosophila* and assessed for the formation of inclusions. We generated flies expressing different phosphomutant versions of 97QHTTex1 fused to mCherry in adult dopaminergic neurons, under the control of TH-GAL4 (Fig 5, columns 1-3), or in larval imaginal discs using the eye-specific GMR-GAL4 driver (S4 Fig). Both T3A (Fig 5B) and T3D (Fig 5E) mutants formed fewer inclusions than non-mutated 97QHTTex1 (Fig 5A), while S13D (Fig 5F) and S16D (Fig 5G) mutants showed a significantly larger number of aggregates (quantification in Fig 5O). S13A (Fig 5C) and S16A (Fig 5D) showed no difference in the number of inclusions in comparison with non-mutated 97QHTTex1 (Fig 5O). In the larval eye imaginal discs, all phosphomutants showed decreased aggregation, with the exception of S13D mutant that formed more inclusions than the non-mutated 97QHTTex1 (S4 Fig).

**Figure 5.**
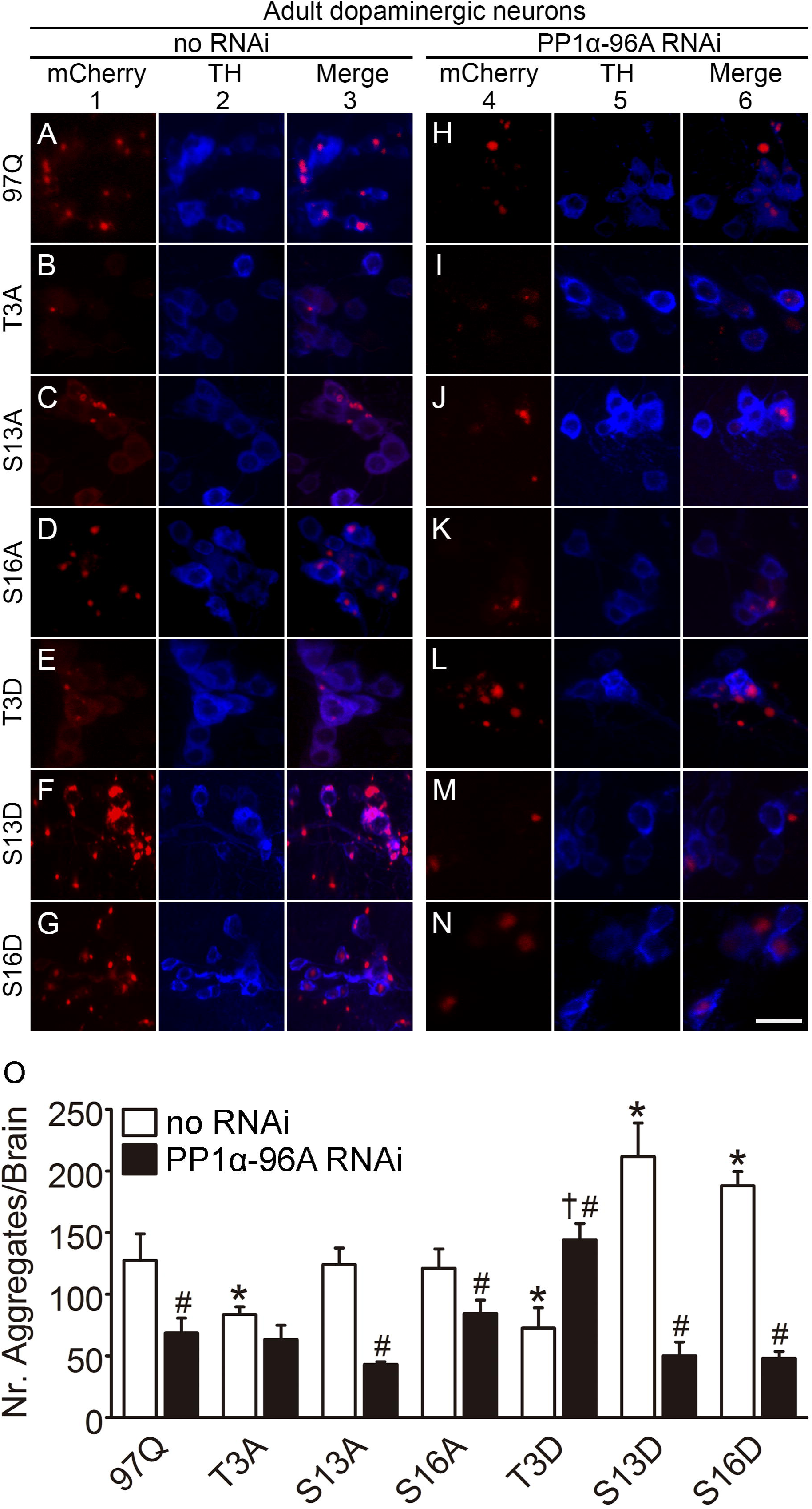
PP1 knockdown prevents 97QHTTex1 aggregation in Drosophila. (A-N) Confocal microscopy images of dopaminergic neurons in adult brains of transgenic flies expressing different versions of 97QHTTex1-mCherry under the control of TH-GAL4. 97HTTex1-mCherry is shown in red (column 1 and 4), and dopaminergic cells (column 2 and 5, anti-TH) are shown in blue. Scale bar, 10 μm. (A-G) *Drosophila* brains expressing non-mutated 97QHTTex1-mCherry or the different phosphomutants. T3A (B) and T3D (E) prevented 97HTTex1-mCherry aggregation, while phosphomimic mutations at S13 or S16 produced a significant increase of 97HTTex1-mCherry inclusion bodies (F and G). (H-N) Co-expression of *PP1α-96A* RNAi together with 97QHTTex1-mCherry or phosphomutants. *PP1α-96A* RNAi decreased the number of 97HTTex1-mCherry inclusions, with the exception of T3A (I), where no change was observed, and T3D (L) expressing flies, which showed a significant increase in 97HTTex1-mCherry aggregation, all in comparison to the respective no RNAi expressing genotype. (O) Quantification of average number of aggregates (±SEM) from at least 5 *Drosophila* brains. *, significant versus TH-GAL4>97QHTTex1. #, significant versus their “no RNAi” transgenic control (white bars). †, significant versus TH-GAL4>97QHTT/*PP1α-96A* RNAi, p<0.05 (one-way ANOVA).

These findings indicate that expression of N17 phosphomutants can modulate 97QHTTex1 aggregation in *Drosophila*, depending on the developmental stage and cellular context.

### PP1 regulates HTTex1 aggregation and neurotoxicity in *Drosophila*

In order to test if protein phosphatases also regulate HTTex1 aggregation in *Drosophila*, we performed RNAi knockdown experiments for the homologues of PP1, PP2A and Cdc25 in flies expressing 97QHTTex1 in dopaminergic neurons (Fig 5, columns 4-6, and S5 Fig). These three phosphatases caused the stronger decrease in HTTex1 aggregation in mammalian cells, upon chemical inhibition. PP1 knockdown flies showed a significant decrease in 97QHTTex1 aggregation (Fig 5H), while PP2A or string (Cdc25 homologue) downregulation had no effect on 97QHTTex1 aggregation (S5 Fig, A and B). PP1 downregulation prevented 97QHTTex1 aggregation in the presence of serine-phosphoresistant mutants (S13A or S16A) (Fig 5J and K), but not T3A, where we did not observe any statistically significant change in the number of aggregates when compared with T3A in the absence of PP1 RNAi (Fig 5O). These results indicate that the effect of PP1 RNAi on 97QHTTex1 aggregation might be mostly mediated by T3 phosphorylation. On the other hand, PP1 knockdown caused an increased number of aggregates in T3D background, and a reduction in the number of aggregates in S13D and S16D backgrounds (Fig 5O). Since HTTex1 does not contain any phosphorylatable residue beyond N17 domain, the increased aggregation observed in the T3D background upon PP1 inhibition suggests that S13 and/or S16 might also be target for phosphorylation to modulate 97QHTTex1 aggregation, in our *Drosophila* model.

We next analysed the effect of PP1, PP2A or string RNAi knockdown on HTTex1 toxicity in the *Drosophila* eye photoreceptor neurons (Fig 6 and S5 Fig). Importantly, genetic inhibition of PP1 did not compromise rhabdomere viability in flies expressing wild-type (19Q) HTTex1 under the control of Rh1-GAL4 (Fig 6A, column 4 and 6D). However, PP1 knockdown significantly enhanced 97QHTTex1 toxicity, further reducing the number of rhabdomeres in an age dependent manner (Fig 6A-C, column 5 and 6C). PP2A or string RNAi had no effect in the number of rhabdomeres upon co-expression with 19QHTTex1 or 97QHTTex1 (S5 Fig, C and D).

**Figure 6.**
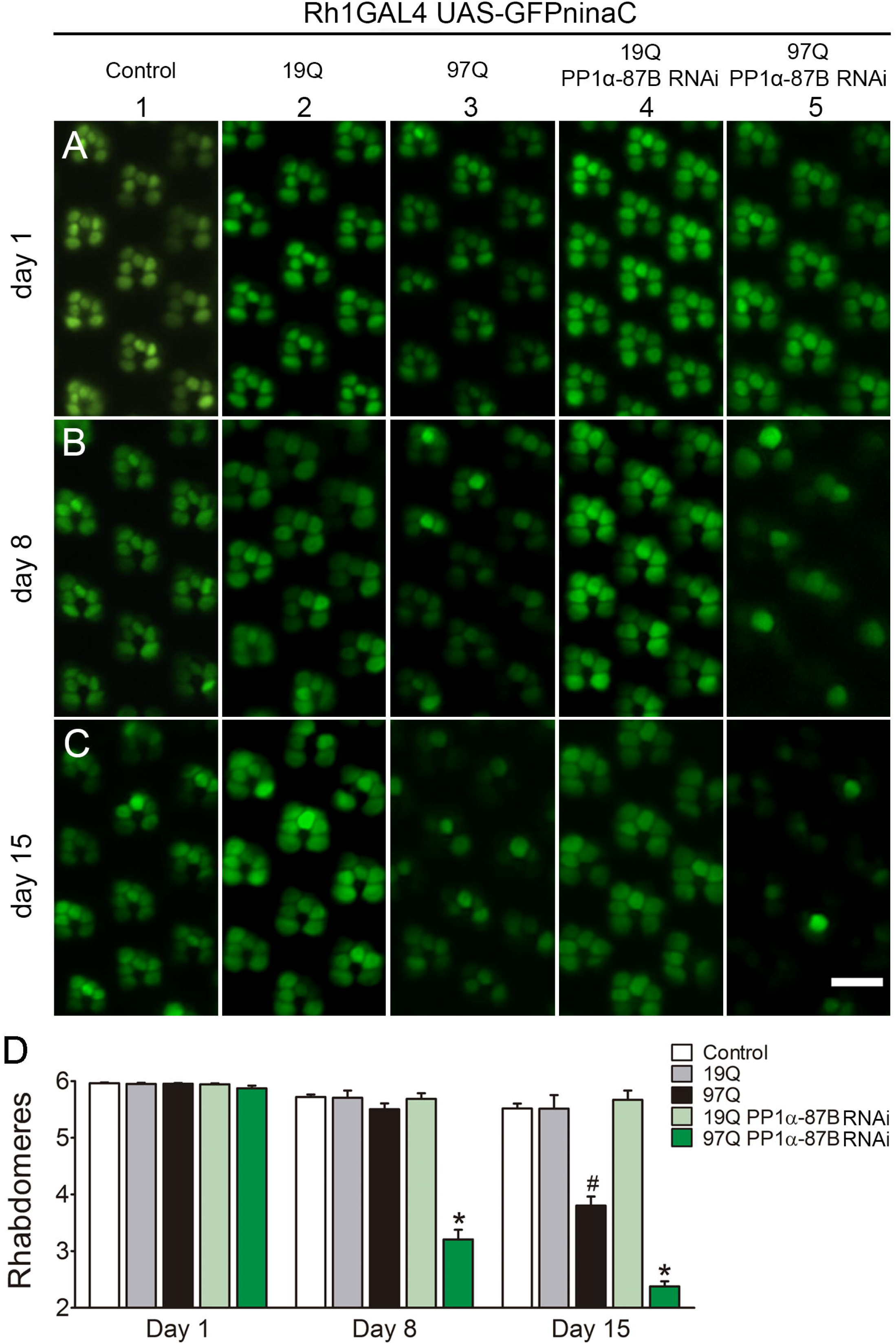
Downregulation of PP1 potentiates HTTex1 neurotoxicity in adult fly photoreceptor neurons. (A-C) Representative pictures of photoreceptors observed by water immersion live imaging of the retinas. Wild-type flies, HD flies and flies expressing *PP1α-87B* RNAi together with 19QHTTex1 or 97QHTTex1 were analysed at day 1, 8 and 15 post-eclosion. Wild-type and 19QHTTex1 expressing flies showed normal retinal morphology as the 6 outer photoreceptors were visible per ommatidium (columns 1 and 2). Flies expressing 97QHTTex1 exhibited age-dependent neurodegeneration with progressive loss of rhabdomeres from 8 to 15 days after eclosion (B and C, column 3). *PP1α-87B* downregulation caused an increase of neurotoxicity in flies expressing 97QHTTex1 (B and C, column 5) and did not affect photoreceptor integrity of 19QHTTex1 flies (column 4). Visualization of the rhabdomeres was done using Rh1-Gal4>UAS-GFP^ninaC^, as described in the methods section. Scale bar, 10 μm. (D) Quantification of mean rhabdomeres (±SEM) per ommatidium. At least 30 ommatidia were analysed from eyes of 5-11 different flies. *, significant versus 97Q. #, significant versus 97Q at day 1 and day 8, p<0.05 (two-way ANOVA).

## Discussion

Protein misfolding and aggregation are intimately involved in the pathogenesis of HD and other neurodegenerative disorders. Despite extensive research in the field, the precise molecular mechanisms by which misfolded proteins aggregate and form toxic species remain elusive. Increasing evidence suggests that targeting phosphorylation events in the N17 domain of mutant HTT can influence the pathological function of the protein [28,30,36–38]. However, the significance of single N17 phosphorylation events in HTT oligomerization, aggregation and toxicity is still poorly understood.

Here, we report that single N17 phosphorylation can prevent HTTex1 aggregation, but not oligomerization. We propose that each residue has a different “strength” in modulating HTTex1 aggregation. Importantly, we found that protein phosphatase 1 can control HTTex1 aggregation and toxicity, suggesting that N17 phosphorylation might be mediated by this protein phosphatase. The fact that single N17 phosphorylation events are sufficient to abolish HTTex1 aggregation could be very important from an HD therapeutic perspective, since a single N17 phosphorylation could provide a simpler molecular target than double phosphorylation.

We show that single phosphomimic mutation in T3, S13 or S16 allow 97QHTTex1 to oligomerize but not to form large inclusions in human cells (Fig 1 and S2 Fig). Double S13/S16 phosphorylation prevents aggregation both *in vitro* and *in vivo*, and toxicity *in vivo* [22,26]. However, double S13/S16 phosphorylation is less abundant than single phosphorylation events [39], and may require the overexpression of specific kinases [30], which may confound therapeutic efforts based on this approach. Although some studies reported on the effect of single S13 or S16 phosphorylation events on HTTex1 intracellular localization, they did not focus on HTTex1 oligomerization, aggregation or toxicity [28,37]. Results from cell-free systems indicate that single S13- or S16-phosphorylated HTTex1 behave similarly to double S13/S16-phosphorylated HTTex1 in terms of aggregation, being unable to form mature fibrils [26]. Our findings in living human cells provide additional biological support for those *in vitro* studies.

The unique properties of our BiFC cellular model allowed us to analyse the relative contribution of each phosphorylatable residue towards HTTex1 oligomerization and aggregation. We found that T3D completely abolished 97QHTTex1 aggregation in the presence of any other phosphomutant or non-mutated 97QHTTex1 molecules, while S13D or S16D only had a partial reduction effect (Fig 2 and Table 1). These results, together with the higher abundance of T3-phosphorylated pools [39,40], highlight the relevance of this residue in modulating the aggregation of HTTex1, and strongly support T3 as a promising target for HD intervention [29]. Our results also suggest that, HTTex1 might be predominantly unphosphorylated under pathological conditions [29,37], as phosphoresistant combinations behave more similarly to the non-mutated 97QHTTex1 pair than to the phosphomimics, regarding their aggregation pattern (Fig 1 and Table 1).

Previous *in vitro* studies demonstrate that N17 phosphorylation inhibits β-sheet conformation of mutant HTTex1 and suppresses its fibrillisation, stabilizing the α-helical structure of the N17 domain which could led to altered aggregation dynamics [41,53]. In FRAP experiments, we show that combinations of phosphomimic and non-mutated 97QHTTex1 induced the formation of inclusions that were more dynamic, diffuse and fluid than regular 97QHTTex1 aggregates, resembling an intermediate stage between oligomers and mature inclusions (Fig 3, S1-S4 and S6 Videos). Additionally, non-mutated 97QHTTex1 pairs did not aggregate to the same extent as the phosphoresistant combinations (Fig 1C and Table 1). Thus, we propose a model where unphosphorylated HTTex1 fragments oligomerize and form inclusions that grow into mature fibrils until enough phosphorylated HTTex1 molecules are intercalated in the structure to interfere with the process, acting as a ‘brake’. This could explain the existence of aggregates of various sizes and morphologies.

Genetic and pharmacological inhibition of PP1 resulted in lower 97QHTTex1 aggregation and increased toxicity (Fig 4-6). Protein phosphatases PP2B/3 (Calcineurin), PP2C and PP1/PP2A have been also shown to regulate HTT phosphorylation at several residues beyond exon 1, as well as HTT toxicity [42–46]. It is important to note that HTTex1 does not contain any phosphorylatable residue beyond the N17 domain and, therefore, any direct effect of protein phosphatases should happen on T3, S13 or S16. Our results indicate that PP1 affects 97QHTTex1 aggregation by regulating T3 phosphorylation. In *Drosophila* dopaminergic neurons, PP1 knockdown in serine-phosphomutant backgrounds leads to a decrease in 97QHTTex1 aggregation, regardless the phosphorylation-like state of these residues (Fig 5O). Moreover, PP1 RNAi does not cause any further reduction of 97QHTTex1 aggregation when co-expressed with T3A mutant. Together, these data suggest that the effect of PP1 inhibition on HTTex1 aggregation is primarily mediated by T3 phosphorylation. However, an increase in 97QHTTex1 aggregation is observed when PP1 RNAi is co-expressed with T3D mutant (Fig 5L and O), indicating that S13 and S16 might also be target for PP1 regulation. Since S13D or S16D increase 97QHTTex1 aggregation (Fig 5F, G and O), we hypothesize that PP1 modulates S13 or S16 phosphorylation upon T3 phosphorylation. In fact, it is likely that different N17 mutations may contribute to subsequent phosphorylation events in other residues. For example, IKK-mediated S16 phosphorylation is facilitated by previous phosphorylation of S13 [30].

Interestingly, the striking decrease in 97QHTTex1 aggregation observed when PP1 RNAi was co-expressed with S13D or S16D mutants versus S13D or S16D alone (no RNAi) (Fig 5O), suggests that T3 phosphorylation is dominant over S13 or S16 phosphorylation. Our human cell data also supports this hypothesis, since S13D or S16D BiFC combinations with non-mutated 97QHTTex1 still shows aggregates, while T3D BiFC combinations with any other mutant completely abolishes aggregation (Fig 2 and Table 1).

In summary, our results support a strong role for single N17 phosphorylation events on HTTex1 aggregation, dynamics and toxicity, and uncover the regulatory role of PP1 in these events. Ultimately, our study opens novel avenues for the therapeutic targeting of PP1 and N17 phosphorylation in HD.

## Materials and methods

### Cell culture, plasmids and treatments

Human H4 glioma cells (ATCC HTB-148, LGC Standards, Barcelona, Spain) were maintained in OPTI-MEM I culture medium (Gibco, Invitrogen, Barcelona, Spain) supplemented with 10% (v/v) fetal bovine serum (FBS) and 1% (w/v) of a penicillin/streptomycin commercial antibiotic mixture (Gibco, Invitrogen, Barcelona, Spain), under controlled conditions of temperature and humidity (37°C, 5% CO_2_). Different types of cell culture dishes were used for cell seeding depending on the application. For flow cytometry and toxicity assays, cells were grown on 6-well plates (35 mm diameter, Techno Plastic Cultures AG, Switzerland). For microscopy, cells were seeded on glass-bottom 35 mm dishes (10 mm glass surface diameter, MatTek Corporation, Ashland, MA, USA). And, for protein extraction (PAGE and filter trap assays), cells were seeded on 100 mm dishes (Techno Plastic Cultures AG, Switzerland). For all experiments, cells were counted and seeded at a density of 10,000 cells/cm^2^ regardless dish size. Generation of HTTex1- and tau-Venus BiFC constructs was described in detail elsewhere [47,49]. Alpha-synuclein-Venus BiFC plasmids were a kind gift from Pamela J. McLean (Department of Neurology, Alzheimer’s Disease Research Unit, Massachusetts General Hospital, MA, USA). Phosphomimic (T3, S13 or S16 mutated to aspartic acid) and phosphoresistant (T3, S13 or S16 mutated to alanine) constructs were produced by PCR-based site-directed mutagenesis using 97QHTTex1-Venus plasmids as templates. Plasmid transfection was performed by means of the X-tremeGene 9 reagent (Roche diagnostics, Mannheim, Germany), following manufacturer’s instructions. Twenty-four hours after transfection, cells were collected and analysed for oligomerization, aggregation and toxicity as described below. Pharmacological inhibition of protein phosphatases was performed using a phosphatase inhibitor library (Enzo Life Sciences, Lausen, Switzerland). Briefly, cells were treated with 33 different phosphatase inhibitors upon transfection of 97QHTTex1 BiFC constructs. Phosphatase inhibitors were dissolved in DMSO and added to culture medium at variable concentrations, according to the IC50 described in manufacturer’s instructions (S1 Table).

### Flow cytometry

Cells were washed with Ca^2+^ and Mg^2+^ free phosphate buffer saline (PBS) (Gibco, Invitrogen, Barcelona, Spain) and collected by trypsinization (0.05% w/v trypsin, 5 min, 37°C) into BD Falcon Round-Bottom tubes (BD Biosciences, San Jose, CA, USA). Cell pellet was resuspended in PBS and analysed by means of a LSR Fortessa flow cytometer (Beckton Dickinson, Franklin Lakes, NJ, USA). Ten thousand cells were examined per experimental group. The FlowJo software (Tree Star Inc., Ashland, OR, USA) was used for data analyses and representation.

### Fluorescence microscopy and FRAP experiments

Images of living H4 cells were acquired using an Axiovert 200M widefield fluorescence microscope equipped with a CCD camera (Carl Zeiss MicroImaging GmbH, Germany). Pictures of a total of 100-150 cells per sample were scored for aggregate quantification using the ImageJ free software (http://rsbweb.nih.gov/ij/). FRAP experiments were performed using a META LSM 510 confocal microscope. Briefly, protein aggregates were focused at the central focal plane and adjusted to avoid pixel saturation. Experiments lasted for 70-150 s, taking one picture every second. After establishing the basal signal, aggregates were bleached using the 488 nm laser line at 100% laser transmission on a circular region of interest (ROI) with a diameter of 30 pixels (1.31 μm radius) for 5 s (10 iterations). Fluorescence recovery was then monitored for 60-140 s with LSM software. Images were analysed and prepared for publication by means of the ImageJ free software.

### Immunoblotting

Proteins were extracted in native or denaturing conditions according to the requirements of each technique. Briefly, cells were washed with PBS 1X and collected by scraping. Cells were incubated with lysis buffer and sonicated for 10 sec at 5 mA using a Soniprep 150 sonicator (Albra, Milano, Italy). For denaturalizing conditions, the lysis buffer was 1% Triton X-100, 150 mM NaCl, 50 mM Tris pH 7.4, supplemented with a protease inhibitor cocktail (Roche diagnostics, Mannheim, Germany). For native conditions, the lysis buffer was 173 mM NaCl, 50 mM Tris pH 7.4, 5 mM EDTA, also supplemented with protease inhibitor cocktail. Proteins were collected after cell lysate centrifugation at 10,000 g for 10 min at 4°C and quantified by means of the BCA Protein Assay Reagent Kit (Thermo Fisher Scientific Inc., Rockford, IL, USA), following manufacturer’s instructions. For SDS- or Native-PAGE immunoblotting, 15 μg of total protein extracts were prepared and separated by electrophoresis using a 12% SDS-polyacrylamide gel or a 5% SDS-free-polyacrylamide gel, respectively. For denaturing conditions, samples were boiled in standard loading buffer (200 mM Tris-HCl pH 6.8, 8% SDS, 40% glycerol, 6.3% β-mercaptoethanol, 0.4% bromophenol blue) for 5 min at 95°C. For native conditions, extracts were mixed with SDS- and mercaptoethanol free loading buffer (200 mM Tris-HCl pH 6.8; 40% glycerol; 0.4% bromophenol blue) and the boiling step was omitted. Proteins were transferred onto PVDF membranes and blocked with 5% (w/v) non-fat dry milk in Tris-HCl buffer saline-Tween solution (TBS-T) (150 mM NaCl, 50 mM Tris pH 7.4, 0.5% Tween-20) for 1 h at room temperature. Membranes were incubated with primary antibodies against HTT (1:500, Millipore, Billerica, MA, USA) and GAPDH (1:30000, Ambion, Austin, TX, USA) as specified. A secondary mouse IgG Horseradish Peroxidase-linked antibody (1:10000, GE Healthcare Life Sciences, Uppsala, Sweden) was used for 1 h incubation at room temperature. Immunoblots were developed with enhanced chemiluminescence reagents (Millipore, Billerica, MA, USA) and exposed to X-ray films.

### Filter trap assays

Cells were pelleted by centrifugation at 700 g for 10 min and cell lysates collected in native conditions as described above. One hundred μg of native protein extracts were mixed with SDS to a final concentration of 0.4% (w/v). Samples were loaded on a dot-blotting device and filtered by vacuum through cellulose acetate membranes (0.22 μm pore; GE Water & Process Technologies, Fairfield, CT, USA), previously incubated with 1% (w/v) SDS solution in PBS. After filtration, membranes were washed twice and processed for immublotting detection of HTT, as described above. In these conditions, only large SDS-insoluble aggregates are retained in the filter and therefore HTT signal is proportional to the presence of large insoluble species. Analyses and quantification of blots signal were performed using ImageJ software. HTTex1 aggregation levels (Fig 4) were calculated by densitometry analyses and normalized to total HTT expression levels and GAPDH loading control.

### *Drosophila* stocks, genetics and crosses

Flies were maintained at 25°C and raised on standard cornmeal medium in a light/dark cycle of 12 h. We generated eight constructs encoding for different versions of HTTex1 fused to mCherry: a wild-type version with a polyQ tail containing 19 glutamines, a mutant version with 97 glutamines, and six constructs encoding phosphomutant versions (T3A/D, S13A/D and S16A/D) with 97 glutamines. To establish transgenic UAS-HTTex1-mCherry lines, our constructs were cloned into pWalium10-roe by the Gateway cloning technology (Thermo Fisher Scientific, USA), and then injected into *y*^*1*^*w*^*1118*^ embryos using phiC31 integrase-mediated DNA recombination (BestGene strain #9723, attP landing site at 2L-28E7). For genetic knockdown experiments in *Drosophila*, we employed UAS-RNAi-targeted lines of four phosphatases: *string* (homolog of cdc25 phosphatase, BL34831, HMS00146), *PP1α-96A* (alpha-1 isoform of PP1 catalytic subunit, BL42641, HMS02477), *PP1α-87B* (alpha-2 isoform of PP1 catalytic subunit, BL32414, HMS00409), and *PP2A-29B* (PP2A regulatory subunit, BL43283, HMS01921). RNAi stocks were obtained from the TRiP (Transgenic RNAi Project) library [54,55], courtesy of the Bloomington Drosophila Stock Center (Indiana University, Bloomington, IN, USA). Three different driver lines were used: TH-GAL4 (active in dopaminergic neurons, under the control of the tyrosine hydroxylase promotor), GMR-GAL4 (active in the eye) and Rh1-GAL4 (active in the photoreceptors R1-R6, under the control of rhodopsin1 promotor). To analyze the effect of RNAi downregulation of specific phosphatases upon HTTex1 aggregation and toxicity, UAS-97QHTTmCherry/CyO;THGAL4/TM6B was crossed with UAS-RNAi-targeted lines. Adult flies carrying UAS-97QHTTmCherry/+;TH-GAL4/UAS-*string*RNAi, UAS-97QHTTmCherry/+;TH-GAL4/UAS-*PP1α-87B*RNAi, UAS-97QHTTmCherry/UAS-*PP1α-96A*;TH-GAL4/+ and UAS-97QHTTmCherry/+;TH-GAL4/UAS-*PP2A-29B*RNAi were selected and analysed. Unless specified otherwise, flies carrying UAS-lacZ targeted to the same location were used as controls. For deep pseudopupil (DPP) analysis, Rh1-GAL4;UAS-GFP^ninaC^/UAS-19QHTTmCherry;UAS-*PP1-87B*RNAi/+ and Rh1-GAL4;UAS-GFP^ninaC^/UAS-97QHTTmCherry;UAS-*PP1-87B*RNAi/+ adult flies were selected and analysed. Rh1-GAL4;UAS-GFP^ninaC^, Rh1-GAL4;UAS-GFP^ninaC^/UAS-19QHTTmCherry;UAS-GFP and Rh1-GAL4;UAS-GFP^ninaC^/UAS-97QHTTmCherry;UAS-GFP were used as controls.

### Immunohistochemistry and confocal microscopy

For confocal imaging of adult brains, 10-days-old flies were dissected and brains prepared as previously described [51]. Briefly, adult flies were anesthetized with CO_2_ and brains isolated in PBS 1X from the head cuticles before being fixed in 4% paraformaldehyde-containing PBS. Dopaminergic neurons were stained by incubation for 48 h at 4°C with mouse anti-TH antibody (1:100, Immunostar, Hudson, WI, USA) in PBST (1X PBS, 0.3% Triton X-100) containing 5% (v/v) normal goat serum. Samples were washed three times for 15 min in PBST. Mouse anti-Cy5 (1:200, Jackson ImmunoResearch, West Grove, PA, USA) was diluted in PBST-containing 5% (v/v) normal goat serum and used as secondary antibody by incubation for 24 h at 4°C. For larva immunohistochemistry, eye imaginal discs of 3^rd^ instar larvae were dissected in PBS 1X and fixed for 1 h in 4% paraformaldehyde as described [56]. Samples were then washed three times for 15 min in PBST and incubated overnight at 4°C with rat anti-Elav antibody (1:100, Developmental Studies Hybridoma Bank, University of Iowa, USA). Imaginal discs were washed three times for 15 min in PBST and incubated for 2 h with rat anti-Cy5 secondary antibody (1:200, Jackson ImmunoResearch, West Grove, PA, USA). Finally, all samples were submitted to a last step of washing and mounted in 80% glycerol PBS solution, followed by confocal microscopy analyses. Z-stack images were acquired using a LSM 710 Meta Zeiss confocal microscope (resolution of 1024 × 1024, slice thickness of 1 μm, frame average of 2). Z-projections were generated and merged using ImageJ free software and images were prepared on Adobe Photoshop CS6 (Adobe Systems Incorporated, San Jose, CA, USA). For quantification of aggregates, mCherry Z-stack images were analysed by means of the Fiji software [57] and aggregation was measured using 3D Objects Counter plugin [58]. The minimum threshold value was defined as 0.144 μm^3^ to exclude signal from soluble HTT in the count. Volume (μm^3^) occupied per each aggregate, average number of aggregates and standard error were calculated for at least 5 imaginal discs or adult brains per genotype.

### Live imaging of adult *Drosophila* eye

DPP analysis was performed in living animals as previously described [51,59]. Briefly, flies at age of 1, 8 or 15 days were anaesthetized with CO_2_ and then placed on a 50 mm petri dish, previously poured with 2% (w/v) agarose at 40°C. Once the agarose was solidified, the anesthetized flies were covered with cool water to keep anesthetic conditions. Adult compound eye integrity of at least 5 flies per genotype was examined by fluorescence microscopy with a water immersion objective (HC APO L40X/0.80W U-V-I, Leica Microsystems, Wetzlar, Germany). Images were obtained using a Leica DM5500 B microscope (Leica Microsystems, Wetzlar, Germany) and an Andor Luca R DL-604M camera (Andor Technology Ltd., Belfast, UK). Images were analysed using Image J free software and number of fluorescing rhabdomeres was scored for >15 ommatidia per fly.

### Statistical analyses

GraphPad Prism 5 (GraphPad Software Inc., La Jolla, CA, USA) or Sigmaplot software (Systat Software, Inc., San Jose, CA, USA) were used to perform the statistical analysis and graphical representation of data. *In vitro* results are shown as the average ± standard deviation (SD) of at least 3 independent experiments and *Drosophila* results as the average ± standard error (SEM), unless specified otherwise. Cell culture data was analysed by means of a one-way ANOVA followed by a post-hoc Tukey test for average comparison. Aggregates quantification in *Drosophila* was analysed by means of a one-way ANOVA with Newman-Keuls post-hoc test. DPP assays were analysed by means of a two-way ANOVA followed by a Bonferroni post-hoc test. Results were in all cases considered significant only when p<0.05.

## Acknowledgements

The authors thank the Bloomington Drosophila Stock Center and the Developmental Studies Hybridoma Bank for fly stocks and antibodies, respectively. The authors also thank Bioimaging Unit from Instituto de Medicina Molecular and Advance Imaging Unit from Gulbenkian Science Institute for support with imaging. TFO and FH were supported by a seed grant from the European Huntington Disease Network (EHDN). TFO is currently supported by the DFG Center for Nanoscale Microscopy and Molecular Physiology of the Brain. FH and PMD were also supported by Project LISBOA-01-0145-FEDER-007660 (Cellular Structural and Molecular Microbiology) funded by FEDER funds through COMPETE2020 - Programa Operacional Competitividade e Internacionalização (POCI) and by national funds through Fundação para a Ciência e Tecnologia (Refs. SFRH/BPD/63530/2009, IF/00094/2013/CP1173/CT0005 and UID/CBQ/04612/2013). JBS was supported by Fundação para a Ciência e a Tecnologia (Ref. SFRH/BD/85275/2012). PMD, GMP and YPA were also supported by Fundação para a Ciência e a Tecnologia (FCT-ANR/NEU-NMC/0006/2013 and PTDC/NEU-NMC/2459/2014). FG thanks the Medical Research Council (MRC) for funding that provided valuable infrastructure supporting this work.

## Author contributions

JBS and FH carried out the experiments in mammalian cells. JBS, GMP and YPA performed the *Drosophila* experiments, under supervision of PMD and FG. FH, JBS, PMD and TFO wrote the manuscript, which was edited by all authors. JBS and FH prepared the figures. FH and TFO had the original idea, supervised the work in mammalian cells and coordinated the different teams.

## Supporting information

**S1 Fig. Single N17 phosphoresistant mutations do not alter HTTex1 aggregation pattern.** Graphic representations from microscopy data of H4 cells transfected with the corresponding HTTex1-Venus BiFC pairs. Single phosphoresistant mutations did not change significantly the number of aggregates per cell (A), the distribution and number of aggregates per cell (B) or the size of aggregates (C).

**S2 Fig. N17 mutations do not affect HTTex1 oligomerization.** (A) Flow cytometry charts of H4 cells transfected with the indicated HTTex1-Venus BiFC pair. Y axis represents the side scatter (SSC) signal, and the X axis represents the signal in the FITC channel. Both scales are logarithmic. (B and C) Graphic representation of quantitative flow cytometry results. *significant versus 19QHTTex1 pair, p<0.05.

**S3 Fig. Combinations of phosphomimic and non-mutated 97QHTTex1 produce similar levels of fluorescence.** (A) Representative flow cytometry profiles of cells co-transfected with combinations of phosphomimic mutants and non-mutated 97QHTTex1 BiFC constructs. (B) Quantitative analysis of flow cytometry data. *significant versus 19QHTTex1 pair, p<0.05.

**S4 Fig. HTTex1 phosphomutants expression in 3^rd^ instar larva eye imaginal discs.** (A-H) Confocal microscopy images of eye imaginal discs in larvae expressing 97QHTTex1-mCherry under the control of GMR-GAL4. 97QHTTex1-mCherry is shown in red, and photoreceptors (anti-Elav) are shown in blue. Mutations in the N17 phosphorylatable residues reduce 97QHTTex1-mCherry aggregation in larvae (C-E and G-F), with the exception of S13D (F), which shows increased number of aggregates versus non-mutated 97QHTTex1 (B). Scale bar, 10 μm.

**S5 Fig. Knockdown of PP2A or *string* does not affect 97QHTTex1 aggregation or toxicity in *Drosophila*. (**A) Imaging of adult dopaminergic neurons in RNAi transgenic flies expressing 97QHTTex1 under the control of TH-GAL4. *PP2A-29B* or *string* RNAi knockdown flies showed no overt phenotype in terms of aggregation when compared to 97QHTTex1 no RNAi control. (B) Quantitative analyses of confocal pictures. Data are average number of aggregates ± SEM. Scale bar, 10 μm. *PP2A-29B* (C) or *string* (D) downregulation did not affect progressive photoreceptor loss observed in 97QHTTex1 flies. *significant versus 97Q, #significant versus 97Q at day 1 and day 8, p<0.05. ns, no significant versus 97Q at day 15.

**S6 Fig. Effect of phosphatase inhibitors on HTTex1 expression.** Representative blots of total protein extracts from H4 cells transfected with non-mutated 97QHTTex1-Venus BiFC constructs and incubated with the indicated phosphatase inhibitors for 24 h. Membranes were probed with anti-HTT and anti-GAPDH antibodies, as indicated. Corresponds to S1 Table. D, DMSO.

**S7 Fig. Full images of the western blot figures presented in this article.** A-C, blots corresponding to Fig 1D. D, blots corresponding to S6 Fig.

**S1 Video. Fluorescence recovery after photobleaching of an aggregate constituted exclusively by non-mutated 97QHTTex1.** Corresponds to the time-lapse shown in Fig 3A, first row (97Q/97Q).

**S2 Video. Fluorescence recovery after photobleaching of an aggregate constituted by combination of non-mutated 97QHTTex1 and S13D.** Corresponds to the time-lapse shown in Fig 3A, second row (S13D/97Q).

**S3 Video. Fluorescence recovery after photobleaching of an aggregate constituted by combination of non-mutated 97QHTTex1 and S16D.** Corresponds to the time-lapse shown in Fig 3A, third row (S16D/97Q).

**S4 Video. Flexibility of intracellular HTTex1 aggregates constituted by combination of non-mutated 97QHTTex1 and S13D.** Corresponds to the time-lapse shown in Fig 3D (S13D/97Q).

**S5 Video. Flexibility of intracellular HTTex1 aggregates constituted by combination of non-mutated 97QHTTex1 and S13A.**

**S6 Video. Flexibility of intracellular HTTex1 aggregates constituted exclusively by non-mutated 97QHTTex1.**

**S1 Table. Name, concentration, description, toxicity and effect on HTTex1 expression of the phosphatase inhibitor library.**

